# A simple, versatile and robust centrifugation-based filtration protocol for the isolation and quantification of α-synuclein monomers, oligomers and fibrils: towards improving experimental reproducibility in α-synuclein research

**DOI:** 10.1101/772160

**Authors:** Senthil T. Kumar, Sonia Donzelli, Anass Chiki, Muhammed Muazzam Kamil Syed, Hilal A. Lashuel

**Affiliations:** Laboratory of Molecular and Chemical Biology of Neurodegeneration, Brain Mind Institute, EPFL, Switzerland

**Author notes:** To whom correspondences should be addressed: Laboratory of Molecular and Chemical Biology of Neurodegeneration, Brain Mind Institute, Ecole Polytechnique Fédérale de Lausanne, 1015 Lausanne. List of abbreviations aSyn- Alpha-Synuclein; PD- Parkinson’s disease; LBs- Lewy bodies; DLB- dementia with Lewy bodies; MSA- multiple system atrophy; CSF- cerebrospinal fluid; EM- electron microscopy; AFM- atomic force microscopy; CD- circular dichroism spectroscopy; SDS- PAGE- sodium dodecyl sulphate-polyacrylamide gel electrophoresis; UV- ultra-violet; BCA- bicinchoninic acid; kDa- kilodalton; MDa- megadalton; TFA- trifluoroacetic acid; WT- wild-type; PBS- phosphate buffered saline; TBS- tris buffered saline; SEC- size exclusion chromatography; nm- nanometer; PFF- pre-formed fibril; MWCO- molecular weight cut-off.

**Keywords:** Alpha-Synuclein, Parkinson’s disease, Amyloid fibrils, Oligomers and Monomers

## Abstract

Increasing evidence suggests that the process of alpha-synuclein (aSyn) aggregation from monomers into amyloid fibrils *via* oligomeric intermediates plays an essential role in the pathogenesis of different synucleinopathies, including Parkinson’s disease (PD), multiple system atrophy and dementia with Lewy bodies. However, the nature of the toxic species and the mechanisms by which they contribute to neurotoxicity and disease progression remain elusive. Over the past two decades, significant efforts and resources have been invested in studies aimed at identifying the putative toxic species along the pathway of aSyn fibrillization, and to develop small molecule drugs or antibodies that target toxic aSyn oligomeric intermediates. Although this approach has helped to advance the field and provide insights into the biological properties and toxicity of different aSyn species, many of the fundamental questions regarding the role of aSyn aggregation in PD remain unanswered, and no therapeutic compounds targeting aSyn oligomers have passed clinical trials. Several factors have contributed to this slow progress, including the complexity of the aggregation pathways and the heterogeneity and dynamic nature of aSyn aggregates. In the majority of experiment, the aSyn samples used contain mixtures of aSyn species that exist in an equilibrium and their ratio changes upon modifying experimental conditions. The failure to quantitatively account for the distribution of different aSyn species in different studies has contributed not only to experimental irreproducibility but also to misinterpretation of results and misdirection of valuable resources. Towards addressing these challenges and improving experimental reproducibility in Parkinson’s research, we describe here a simple centrifugation-based filtration protocol for the isolation, quantification and assessment of the distribution of of aSyn monomers, oligomers and fibrils, in heterogeneous aSyn samples of increasing complexity. The protocol is simple, does not require any special instrumentation and can be performed rapidly on multiple samples using small volumes. Here, we present and discuss several examples that illustrate the applications of this protocol and how it could contribute to improving the reproducibility of experiments aimed at elucidating the structural basis of aSyn aggregation, seeding activity, toxicity and pathology spreading. This protocol is applicable, with slight modifications, to other amyloid-forming proteins.

**Table of Content Figure:** 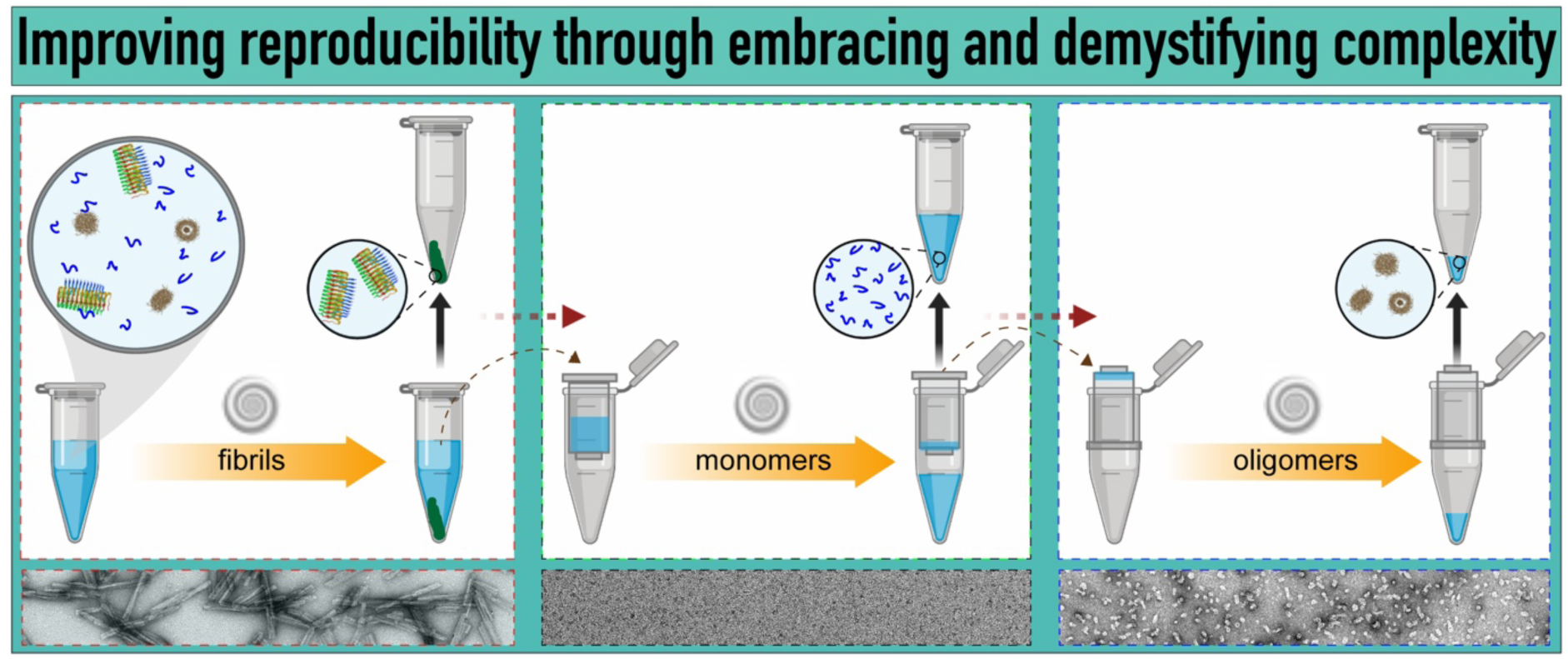

## Introduction

Alpha-synuclein (aSyn) is a 14 kDa protein that is found abundantly in the presynaptic terminals of neurons but is also expressed in smaller amounts in the heart, muscles and other tissues (Jakes *et al*. 1994; Iwai *et al*. 1995). While the precise function of aSyn is not yet well understood, accumulating evidence from genetic and neuropathological studies point to a central role of aSyn in several neurodegenerative disorders that are collectively referred as synucleinopathies (Spillantini & Goedert 2000). These disorders include Parkinson’s disease (PD), dementia with Lewy bodies (DLB), the Lewy body variant of Alzheimer’s disease, multiple system atrophy (MSA) and neurodegeneration with brain iron accumulation (Jellinger 2003; Galvin *et al*. 2001). A common pathological link connecting these disorders is the intracytoplasmic accumulation of misfolded and aggregated forms of aSyn in neurons and in glial cells (Spillantini *et al*. 1997; Baba *et al*. 1998; Takeda *et al*. 1998; Spillantini & Goedert 2000). Despite differences in the cell types and affected brain regions, the fibrillar structures of aSyn enriched in the neuronal inclusions is a salient feature of these diseases (Goedert 2015). aSyn aggregates (oligomers and fibrils) are released by neurons and, upon internalization by the neighbouring cell, induce the misfolding and/or seed the aggregation of endogenous aSyn in the host cells (Danzer *et al*. 2009; Mahul-Mellier *et al*. 2018). This process is thought to be the underlying mechanism for the cell-to-cell spreading and propagation of aSyn to different brain regions during the different stages of disease progression (Hansen *et al*. 2011; Braak *et al*. 2003; Li *et al*. 2008). Although there is a consensus that the process of aSyn fibril formation plays a central role in the initiation and progression of PD and other synucleinopathies, the nature of the toxic species and how they contribute to the different putative disease mechanisms underlying the pathogenesis of PD and other synucleinopathies remain unknown.

Several studies have shown that the presence and/or abundance of Lewy bodies (LBs) do not always correlate with the development of PD symptoms (e.g. the presence of LBs in healthy brains) (Lipkin 1959; Forno 1969; Gibb & Lees 1988; Mikolaenko *et al*. 2005; Duffy & Tennyson 1965; Schmidt *et al*. 1991) or disease severity. These observations, combined with the absence of a direct correlation between aSyn fibril formation and neurodegeneration in different models of PD and the detection of soluble oligomeric forms of aSyn in the cerebrospinal fluid (CSF), have led to a shift of interest from fibrils towards prefibrillar aggregates (oligomers) (Park *et al*. 2011). Therefore, significant efforts and resources have been devoted over the past two decades to dissecting the different steps of aSyn fibril formation *in vitro* and isolating individual species along the pathway of aSyn fibrillization. These efforts led to the development of several protocols for the generation and isolation of aSyn oligomeric species with a distinct secondary structure, size and morphological distribution (Table 2).

Despite many attempts over the past two decades, success has not been achieved in generating homogeneous preparations that comprise a single type of oligomer of a specific size or morphology (Lansbury & Lashuel 2006; Lashuel 2005; Lashuel & Lansbury 2006; Alam *et al*. 2019). Most of the existing methods and protocols led to the generation of a population of oligomers with variable size and morphological distribution that is often in equilibrium with monomers. Depending on the ratio of oligomers to monomers, the propensity for fibril formation in these preparations varies. Very often, chemical cross-linking is used to stabilize oligomers, but this significantly alters their dynamic properties (Nasstrom *et al*. 2011) and the ability to undergo conformational changes that may be required for exerting their biological activity. Therefore, the vast majority of aSyn oligomeric preparations used in research today are heterogeneous in terms of the size distribution of the oligomers and the ratio of monomers, oligomers and fibrils.

During the past 6-8 years, aSyn fibrils have made a striking comeback, mainly due to the emergence of the prion-like hypothesis for the pathological spreading during the progression of PD. This hypothesis suggests that aSyn aggregates (fibrils or oligomers) are secreted by neurons (Lee *et al*. 2005; Lee *et al*. 2008; Desplats *et al*. 2009), and exhibit a potent seeding capacity that allows them to induce aggregation and inclusion formation upon internalization into neighboring neurons and other cell types (Luk *et al*. 2009; Volpicelli-Daley *et al*. 2011; Luna *et al*. 2018; Karpowicz *et al*. 2019). Consistent with this hypothesis, inoculation of fibrils produced *in vitro* (Luk *et al*. 2012; Shimozawa *et al*. 2017; Abdelmotilib *et al*. 2017) or aSyn aggregate-enriched extracts from diseased brains is sufficient to induce Syn fibrillization and the formation of Lewy body-like pathology in neurons, rodent models and non-human primates (Recasens & Dehay 2014; Watts *et al*. 2013; Recasens *et al*. 2014; Masuda-Suzukake *et al*. 2013; Tu *et al*. 1998; Peng *et al*. 2018). This has led to a resurgence in the number of studies focusing on fibrils and strategies to interfere with seeding capacity and role in pathology spreading. The vast majority of this research relies on the ability to reproducibly generate fibril preparations of the desired properties. While incubating concentrated samples of aSyn (3-5 mg/mL) at 37°C with agitation reproducibly leads to the formation of fibrils, very often these preparations contain different amounts of monomers and oligomers. Given that most of these samples are analyzed using electron microscopy (EM) or atomic force microscopy (AFM), which are not quantitative imaging techniques, the presence of the monomers and oligomers is either not detectable (monomers) or under-appreciated because of the extensive amount of fibrils present in the sample. Because of the differences in the dynamic and structural properties of each of the three species, the nature of the samples could change dramatically from one preparation to another and even for the same preparation when exposed to different conditions. Furthermore, all fibril preparations that are used to investigate cellular toxicity, aSyn seeding mechanisms and pathology spreading in cellular and animal models of PD are subjected to sonication or mechanical disruption procedures to generate short fibril seeds with a similar size distribution (50-200 nm). This procedure leads to fibril fragmentation resulting in the generation of more fibril ends and an increase in the amount of monomers/oligomers released from the fibrils. The levels of monomers/oligomers in fibril seed preparation have been shown to be a critical determinant of their dynamics, growth and toxicity (Mahul-Mellier *et al*. 2015). In response to the high variability in aSyn fibril preparations and concerns about reproducibility, there is an urgent need to share the best practices, to develop reproducible protocols and methods for generating aSyn fibrils, and to follow and simple methods for quality control, characterization and validation of such preparations (Polinski *et al*. 2018; Patterson *et al*. 2019).

The increasing reliance on aSyn aggregate preparations (fibrils and oligomers) in the development of PD models and their use to investigate the mechanisms linked to aSyn dysfunction and toxicity underscore the critical importance of using high quality and reproducible preparations in such studies. However, experience teaches us that it is nearly impossible to generate oligomer and fibril preparations of identical species, size and morphology distribution. This challenge could be easily addressed by 1) developing protocols and methods that allow the separation and proper quantification of the amount of each of the three species, and 2) providing basic morphological and structural characterizations of the crude mixtures and isolated species by imaging techniques (EM and AFM) and circular dichroism spectroscopy (CD), respectively. This level of characterization would at least enable better interpretation of the structure-activity studies and improve the reproducibility by allowing accurate comparison of data and results across different laboratories.

Previous studies have shown that different preparations of oligomers and fibrils exhibit different levels of toxicity. The presence of monomers in such preparations has been shown to enhance oligomer and fibril growth as well as the toxicity of these species. (Mahul-Mellier *et al*. 2015; Jan *et al*. 2008). These observations underscore the critical importance of using well-characterized preparations of aSyn samples that contain only monomers, oligomers or fibrils, or preparations where the well-defined concentrations of the different species can be measured. To achieve this goal, one needs to develop methods that not only enable the isolation of different species along the aSyn amyloid pathway but also allow for the rapid assessment of the distribution of the different species prior to or during the experiments aimed at assessing their effects. To improve reproducibility and allow for comparison of oligomer preparations and results from different laboratories, it is crucial that such protocols are simple, do not require complicated or expensive setups or instrumentations, and can be performed within hours to allow direct assessment of the samples immediately before or during the experiments, and in response to changes in experimental conditions.

To address these challenges, we present here a straightforward protocol based on centrifugation and filtration procedures, which allow rapid quantitative and qualitative assessment of the different types of aSyn species such as monomers, oligomers and fibrils in samples of different complexities. This protocol is based on years of experience in our group (Fauvet *et al*. 2012; Khalaf *et al*. 2014) and the evidence that it has been successfully applied in aSyn research by other groups (Zhang *et al*. 2013; Kumar *et al*. 2018; Ghosh *et al*. 2015). The flexibility of this method relies on the fact that it can be used to characterize purified aSyn samples or at different time points during aSyn aggregation. The protocol only requires a minimum sample volume (100 μL) and can be implemented using basic laboratory tools. The total running time of the protocol is 3-4 hours, including 1) isolation of all three species (fibrils, monomers and oligomers, 2) assessment of the distribution of three species by sodium dodecyl sulphate-polyacrylamide gel electrophoresis (SDS-PAGE); and 3) concentration determination using ultra-violet (UV) absorbance and/or bicinchoninic acid (BCA) assay. Biophysical characterization of the aSyn species using CD and EM is necessary to allow batch to batch comparision and to improve experimental reproducibility. Below, we present and discuss several examples that illustrate the potential applications of this protocol, how it could contribute to improving the reproducibility of experiments aimed at elucidating the structural basis of aSyn aggregation, seeding activity, toxicity and pathological spreading.

## Materials and Methods

Key equipment and materials.

**Key equipment and materials**
| **Equipment/material** | **Company** | **Catalogue** |
| --- | --- | --- |
| <u>Centrifuges</u><br>Benchtop | Eppendorf 5417R |  |
| Ultracentrifuge | Beckmann Coulter Optima Max XD |  |
| <u>Spin filters</u><br>30 kDa | Microcon-30 Ultracel YM 30 | MRCF0R030 |
| 50 kDa | Amicon Ultra-2 mL, Ultracel-50K | UFC205024 |
| 100 kDa | Microcon DNA fast flow Ultracel | MRCF0R100 |
|  | Vivaspin 500 | 28-9322-37 |
| Sonicator | Sonics Vibra-Cell™ | VCX 130<br>630-0422 (microtip) |

### Recombinant overexpression and purification of wild type aSyn and cysteine variants

BL21 (DE3) cells transformed with the pT7-7 plasmid encoding the WT aSyn or M1C or A140C aSyn mutants were grown in LB medium at 37 °C and induced with 1 mM 1-thio-β-D-galactopyranoside (AppliChem) at an O.D. between 0.4-0.6 and continued to grow for 4 hours at 37 °C. Induced bacterial cultures were pelleted and sonicated for the lysis of the cells. After centrifugation at 18000*g* for 20 min, the supernatant was boiled for 5 min and centrifuged again for 20 min. The supernatant was purified by anion exchange chromatography (HiPrep 16/10 Q FF, GE Healthcare Life Sciences), followed by reverse-phase HPLC (Jupiter 300 C4, 20 mm I.D. × 250 mm, 10 µm average bead diameter, Phenomenex) and lyophilized. The M1C aSyn mutant was purifed as thiazolidine adducts from bacterial expression. To make cysteine residue available for the formation of disulfide-linked homodimer, purified M1C aSyn was dissolved in reaction buffer (0.1% TFA, 5% acetonitrile, 5 % acetic acid in water) to a final concentration of 0.5 mM. To this solution 100 eq of Silver trifluoromethanesulfonate (Sigma) from 1M solution in 0.1% TFA H_2_O was added. The solution was then incubated at RT without shaking. After 30 min, the reaction was analyzed by electron spray ionization mass spectrometry which showed quantitative deprotection of thiazolidine adducts. The reaction mixture was desalted using PD-10 column using the reaction buffer (GE Healthcare) and lyophilized.

### Preparation WT aSyn oligomers

To prepare WT aSyn oligomers, 60 mg of lyophilized protein was dissolved in 5 mL of phosphate-buffered saline (PBS) (10 mM disodium hydrogenphosphate, 2 mM potassium dihydrogen phosphate (pH 7.4), 137 mM NaCl and 2.7 mM potassium chloride) and followed by incubation for 5 hours at 900 rpm constant shaking at 37 °C. Then, the solution was centrifuged to remove any insoluble particles at 12000*g* for 10 minutes at 4 °C. 5 mL of supernatant was loaded onto a HiLoad^®^ 26/600 Superdex^®^ 200 preparation grade (GE lifesciences) XK prepacked column equilibrated with PBS, and protein was eluted as 2.5 mL fractions at a flow-rate of 1 mL/minute. Fractions corresponding to the void volume peak (oligomers) were split to 500 μL aliquots, snap-frozen and stored at -20 °C.

### Step-by-step explanation of the centrifugation-based filtration protocol

This protocol can be applied to the characterization of aSyn preparations containing variable amounts of aSyn monomers, oligomers or fibrils, such as the freshly dissolved purified protein, oligomers or fibrils. The protocol has been divided into three steps: 1) a centrifugation step for the isolation of the fibrils from the soluble species; 2) a filtration step for the separation of monomers and oligomers; and 3) recovery of the soluble oligomers. A standard procedure for the application of this protocol is described below:

1. A total sample volume between 50 μL to 500 μL is subjected to ultracentrifugation at 100,000*g* for 30 min at 4 °C.
2. After centrifugation, a careful pipetting of the supernatant from the pellet is carried out to isolate the soluble species. The isolated pellets can be used for further analysis after resuspension in the buffer of interest.
3. Next, the supernatant is transferred into the filtration membrane unit (100 kDa), placed into a new collection tube and centrifuged at 15300*g* for 20 min at 4 °C.
4. After centrifugation, the membrane unit is separated from the collection tube. The sample which passes through the membrane and enters the new tube, is named as filtrate sample and contains predominantly monomeric or dimeric aSyn species.
5. To collect the oligomeric species that were retained on the membrane unit, named retentate, 9 parts of the buffer from the initial volume of the sample is added to the membrane (for example, use 450 μL of buffer if 500 μL was the initial volume of the sample), pipette gently 2-3 times and place the filtration membrane unit into a new collection tube by carefully turning it upside-down. Next, pipette one part of the buffer (e.g. 50 μL) into the inverted bottom and centrifuge at low speed at 425*g* for 4 min to recover the retentate sample.

The centrifugal filter membranes were equilibrated with the sample buffer for a short centrifuge run prior to subjecting the samples to filtration

The different aSyn species separated during the centrifugation-based filtration protocol are obtained in standard buffer solutions and thus can be characterized, without the need to exchange buffer, using common protein concentration determination methods, SDS-PAGE analysis, CD spectroscopy, light scattering, sedimentation velocity or electron microscopy, among other techniques.

### Filtration analysis of WT aSyn monomers using different MWCO filter membranes

WT aSyn was dissolved in 600 μL of Tris buffered saline (TBS) pH 7.5 to give a final concentration of 35 μM while placing it on ice cold condition. The solution was ultracentrifuged for 30 min at 100000*g* at 4 °C to remove any preformed insoluble aggregates. The supernatant was collected and filtered through different MWCO centrifugal filters (30 kDa, 50 kDa or 100 kDa). 100 µLwere used for each filtration. The sources of the centrifugal filter membranes used are described in the table above.

For 30 kDa and 100 kDa filtration, 100 μL of the supernatant solution was centrifuged at 15300*g* for 20 min and the filtrate was collected. The retentate was collected by inverted centrifugation using 40 µl of TBS (pH 7.5) at 425 g for 4 min at 4 °C. The volume is diluted to 100 µl using TBS. For the 50 kDa filter, 100 μL of the supernatant was centrifuged at 4000*g* for 20 min at 4 °C in a swinging bucket. The filtrate was collected and diluted to 100 μL using TBS. Next, 100 μL TBS was added to the membrane, pippeted up and down for few times and collected back into a new Eppendorf tube using pippete as the retentate. All the centrifugal filter membranes were equilibrated with TBS by a short centrifuge run prior to subjecting the samples for filtration.

20 μL of each filtrate and retentate for each membrane was mixed with 20 μL of 2× Laemmli buffer for SDS-PAGE analysis and heated at 95°C for 5 min. The samples were analyzed by SDS-PAGE (1.5 mm thickness) immediately or snap frozen in liquid nitrogen and stored at - 80 °C until further use. 10 μL of the SDS-PAGE samples were loaded onto a gel. The remaining filtrate and retentate for each membrane were snap-frozen in liquid nitrogen and stored at -80 °C.

### Filtration analysis of WT aSyn monomers using 100 kDa membranes of different material

WT aSyn was dissolved in 600 μL of TBS (pH 7.5) to a final concentration of 35 μM while placing it on ice cold condition. The solution was ultracentrifuged for 30 min at 100000*g* at 4 °C to remove any insoluble aggregates. The supernatant was collected and filtered through a 100 kDa centrifugal filter composed of two different membrane materials, 1) a Microcon 100 kDa membrane made of regenerated cellulose, and 2) a Vivaspin 500 (100 kDa) made up of polyethersulfone membrane. As stated earlier in the methods section, 100 μL of WT aSyn supernatant was centrifuged at 15300*g* for 20 min at 4°C. The filtrate was collected and diluted to 100 μL using TBS. The retentate from the Microcon membrane was recovered following the usual procedure. However, for Vivaspin, the retentate was collected by addition of 100 μL TBS to the membrane, pippeted up and down for few times and collected back into a new Eppendorf tube using pippete. The SDS-PAGE samples were prepared and analyzed as described above. SDS-PAGE analysis was performed on a 1 mm thick gel.

### Filtration analysis of M1C and A140C aSyn mutants using 100 kDa filter membrane

M1C aSyn was dissolved in 270 μL of TBS, and A140C aSyn was dissolved in 560 μL of TBS (pH 7.5) to reach a final concentration of 35 μM. Both solutions were kept at RT for 3 d without shaking and analyzed using mass spectrometry for validation of dimer formation. After 3 d, the solutions were centrifuged at 100000*g* for 30 min at 4 °C to remove any insoluble aggregates. The supernatants were collected and subjected to the usual filtration analysis using Microcon 100 kDa as follows. 100 µL of supernatant was centrifuged at 15300*g* for 20 min at 4 °C. The filtrate was collected, and the retentate was collected by inverted centrifugation at 425*g* for 4 min at 4 °C using 40 μL of TBS and the volume diluted to 100 μL using TBS. Fractions were collected at each step and equal volumes of samples were used for SDS-PAGE analysis.

### Transmission electron microscopy

Prior to the application of the sample, Formvar and carbon-coated 200 mesh-containing copper EM grids (Electron Microscopy Sciences) were glow-discharged for 30 seconds at 20 mA using a PELCO easiGlow™ Glow Discharge Cleaning System (TED PELLA, Inc). Subsequently, 5 μL of the sample was placed onto the EM grid for a minute. Then, the sample was carefully blotted using filter paper and air-dried for 30 seconds. Then, the grids were washed three times with ultrapure water and stained with 0.7% (w/v) uranyl formate solution. Grids were examined using a Tecnai Spirit BioTWIN electron microscope. The microscope was equipped with a LaB6 gun operated at an acceleration voltage of 80 kV, and images were captured using a 4K × 4K charge-coupled device camera (FEI Eagle).

### SDS-PAGE analysis

Samples for SDS-PAGE were mixed with 2× Laemmli buffer (4% SDS, 20% glycerol, 0.004% bromphenol blue, 0.125M Tris-Cl, 10% 2-mercaptoethanol pH 6.8) and loaded onto 15% polyacrylamide gels. The gel was run at 180 V for 1 hour in running buffer (25 mM Tris, 192 mM Glycine, 0.1% SDS, pH 8.3), followed by staining with a solution of 25% (v/v) isopropanol, 10% acetic acid (v/v) and 0.05% (w/v) Coomassie brilliant blue R (Applichem) and destaining with boiling distilled water.

### Far-UV circular dichroism (CD) spectroscopy

CD spectra of aSyn samples were loaded in a quartz cuvette with a 1-mm path length and collected using a Jasco J-815 CD spectrophotometer operated at 20 °C within the range of 200– 250 nm. Data acquisition used the following parameters: data pitch, 0.2 nm; bandwidth, 1 nm; scanning speed, 50 nm/min; digital integration time, 2 s. A spectrum of each sample is the average of 10 repeats followed by a binomial approximation.

## Results

### A simple centrifugation-based filtration protocol for the separation of aSyn fibrils, monomers and oligomers based on differences in their size and solubility

Although different methods have been shown to be effective in separating fibrillar aSyn from soluble aSyn species and aSyn monomers or oligomers, including size exclusion chromatography (SEC) or density gradient centrifugation (Lashuel *et al*. 2002; Cremades *et al*. 2012; Conway *et al*. 2000; Gao *et al*. 2015), these methods suffer from the requirement of large sample volumes and/or lead to significant dilution of the sample of interest. Furthermore, many of these methods require at least 1-2 hours of processing time per sample and are thus not amenable to rapid analysis of large number of samples. They also require access to specialized instruments (e.g., fast protein liquid chromatography (FPLC)) and expertise that may not be available in all laboratories.

The protocol described here addresses these limitations and takes advantage of 1) the large differences in the size of the three aSyn species (monomers at 14 kDa, oligomers between 28 kDa–1 MDa and fibrils > 2 MDa), and 2) the difference in solubility of the fibrils compared to the monomers and oligomers. The fibrils, but not the oligomers and monomers, can be sedimented by centrifugation. For samples containing the three forms of aSyn, the first step involves the sedimentation and separation of the fibrils from the monomers and oligomers by ultracentrifugation of the sample at 100000*g* for 30 minutes (scheme 1 (1)). This allows for the separation of the sedimentable fibrils as a pellet from the soluble supernatant, which contains both monomers and oligomers of different sizes (scheme 1 (2)). The supernatant (scheme 1 (3)) is then separated into monomers and mixtures of oligomers by filtration through a 100 kDa molecular weight cut-off (MWCO) filter, which allows only the monomers to pass through (scheme 1 (4)). The oligomers are retained on the filter membrane are then recovered, as shown in scheme 1 (5). Determination of the concentration of each species after isolation, using UV absorbance and/or BCA assay, allows for the robust estimation of the distribution of the different species in the original sample. For a more accurate determination of the concentration of each species, we recommend using amino acid analysis. The requirements for the implementation of this protocol are having access only to: 1) the spin filters (Microcon (100 kDa MWCO, Ireland)); 2) a benchtop centrifuge or an ultracentrifuge; 3) a UV absorbance-based spectrometer or a plate reader; and 4) a gel electrophoresis setup. These are standard pieces of equipment to which research laboratories have easy access. Most importantly, this procedure could be performed on sample volumes ranging from 50 – 500 μL.

### Separation of fibrils from soluble aSyn species (oligomers and monomers)

Due to their simplicity, centrifugation-based sedimentation protocols have emerged as a substitute for SEC-based methods, and are commonly used to study protein aggregation by monitoring and quantifying the amount of sedimentable aggregates and the remaining soluble proteins in the supernatant (Khalaf *et al*. 2014; Fauvet *et al*. 2012). These protocols have proven to be reliable and effective in separating fibrils or sedimentable aggregates from the soluble protein species. However, it is crucial to validate such protocols (through analysis of the supernatant) to ensure that fibrils formed by the protein of interest are sedimentable. In most studies, a relatively short and low centrifugal force of approximately 14000–16000*g* is sufficient to sediment mature, fully-grown insoluble aSyn fibrils (Table 1), but not small fibrils that are <150 nm in length. For example, the pre-formed fibril (PFF) seed preparations that are commonly used in cellular seeding assays and animal models of pathology spreading are usually subjected to sonication prior to use. This procedure is used to induce fibril fragmentation with the aim of obtaining a fibril preparation with an average length of 50-150 nm, which has been shown to facilitate their uptake by cells and give the maximum seeding activity (Tarutani *et al*. 2016; Mahul-Mellier *et al*. 2018; Mahul-Mellier *et al*. 2015). Once fragmented to this size, the fibrils are non-sedimentable at this centrifugal force (Table 1). For such preparations, high centrifugal force at 100000*g* is generally applied to sediment shortened/fragmented fibril seeds from the soluble aSyn species.

**Table 1.** Common centrifugal parameters used to prepare different aSyn species.

| aSyn species | Centrifugal parameters | Rationale | References |
| --- | --- | --- | --- |
| WT, A30P, or A53T monomers | 50 µl aliquot at 16000g for 5 min | To remove insoluble material using a benchtop centrifuge | Conway *et al.* 2000 |
| WT aSyn fibrils | 90-µl aliquot at 40000g for 30 min | To measure the amount of fibril pellet remaining during fibril disassembly | Bousset *et al.* 2013 |
| aSyn 30–110 fibrils | 16000g for 30 min | To monitor fibril disassembly when co-incubated with chaperones | Gao *et al.* 2015 |
| Labeled aSyn fibrils | 15000g for 10 min | To remove the free fluorophores after the labeling of fibrils | Fenyi *et al.* 2018 |
| aSyn strain A and B PFFs | 100000g for 15 min | Seeds were resuspended in D<sub>2</sub>O for structural studies | Guo *et al.* 2013 |
| WT aSyn PFFs | 100000g for 30 min | Sedimentation assay for confirmation of pelletable, active PFFs for cellular assays | Volpicelli-Daley *et al.* 2014 |
| WT aSyn PFFs | 20000g for 10 min | To sediment down PFFs after sonication | Fares *et al.* 2016 |

Taking this into consideration, the protocol described in Scheme 1 can be used to: 1) prepare aSyn monomeric solutions that are free of any pre-formed aggregates which may have formed during the process of storage, lyophilization or resolubilization of aSyn samples, 2) separate the non-fibrillized aSyn species (monomers and oligomers) from fibrils and quantify both species and 3) assess the amount of released monomers or the formation of oligomers due to sonication of the fibrils or treatment with chaperones or small molecule disaggregases (Gao *et al*. 2015; DeSantis *et al*. 2012; Bieschke *et al*. 2010); and 4) enable more comprehensive profiling of the complexity of aSyn samples. These capabilities should enable the assessment of batch to batch variations of aSyn sample preparation, comparison of data from different research groups and improving experimental reproducibility across different laboratories. However, for accurate interpretation of the results using this protocol, it is crucial to verify by EM that this separation protocol works for the protein of interest under the experimental conditions used.

### A method for qualitative and quantitative assessment of batch-to-batch variations in the aSyn fibrils preparations

Here, we provide an overview of the standard methods for the preparation and characterization of aSyn fibril preparations and discuss some of the factors that could influence batch-to-batch variations. We illustrate how the protocol described in Fig. 1 can be used to ensure reproducible preparations of PFF seeds for use in seeding-based aggregation assays (Polinski *et al*. 2018; Tarutani *et al*. 2016) or in cellular and animal studies to investigate aSyn pathology formation and spreading (Luk *et al*. 2012; Luk *et al*. 2009; Volpicelli-Daley *et al*. 2011; Abdelmotilib *et al*. 2017; Shimozawa *et al*. 2017). This is achieved by assessing the following paramaters for each PFF preparation: 1) the percentage of fibril formation; 2) the percentage of remaining soluble aSyn species in each preparation; and 3) the relative stability of the fibrils as determined by the amount of soluble aSyn species released after sonication of the fibrils. The protocol can be also modified to assess the presence and amounts of oligomers in such preparations (see below).

**Fig. 1:**
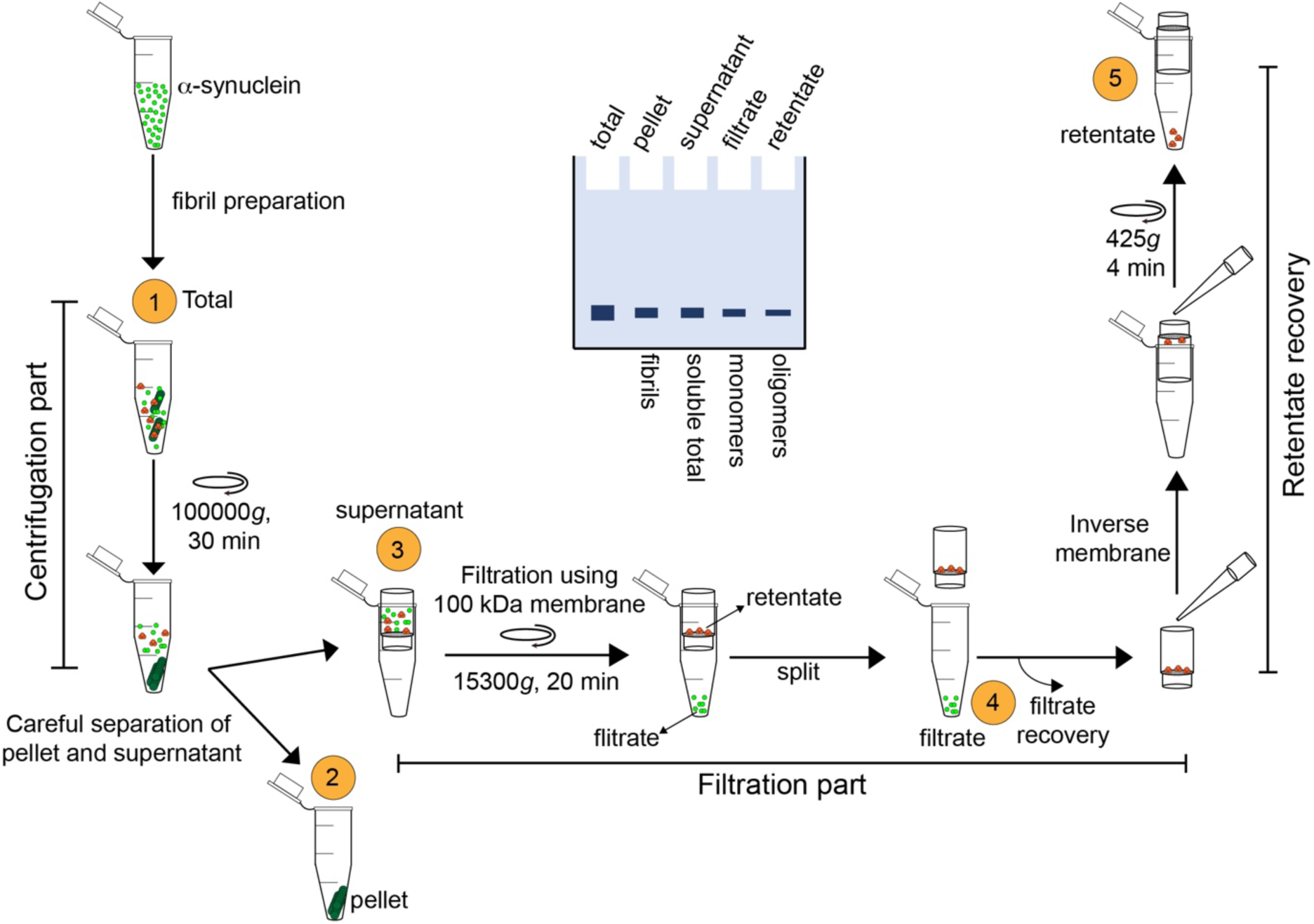
Schematic depiction of the centrifugation-based filtration protocol. Briefly, the protocol includes a centrifugation step for the isolation of the fibril (pellet) from soluble aSyn (monomers and oligomers), a filtration step (through 100 kDa MWCO) for the separation of the monomers from the oligomers, and a final step for the recovery of the oligomers from the spin filters. A depiction of the SDS-PAGE analysis of the samples at each step is shown in the middle. Quantification of band intensity should allow for a quick assessment of the distribution of coexisting aSyn species.

An important criterion for improving the reproducibility of PFF-based experiments is to ensure that the PFF batches are well characterized. The major source of variation between different PFF preparations is the differences in the stability of the fibrils and distribution of aSyn species in the PFF preparations. However, a large concern with the commonly used methods to characterize PFFs is the focus on assessing the morphology of the fibrils and their size distribution rather than focusing on the sample heterogeneity, i.e. the presence of other aSyn species (e.g. monomers and oligomers). Below, we present a series of examples that illustrate protocol described here allows for a more in-depth characterization of the PFF preparations as it also enables the assessment of the levels of aSyn monomers and oligomers species as well as fibril stability, as determined by the extent of fibril-to-monomer disassociation upon PFFs sonication.

For the generation of aSyn fibril preparation, we followed the standard protocols (Mahul-Mellier *et al*. 2018) in which the lyophilized aSyn protein (4 mg/600 μL) was dissolved in Tris·HCl buffer at pH 7.2-7.4 (0.22 μm filtered) and incubated at 37 °C under constant agitation at 1000 rpm on an orbital shaker for 5 days (Fig. 2A). Our experience shows that despite following the same method carefully, it is not unusual to observe variations between different fibrils batches with respect to the extent of the amount of monomers converted to fibrils (Fig. 2 A, C and D) and/or the stability of the fibrils. This is indeed the case for all aSyn fibrillization protocols. Thus, we strongly recommend quantitative assessment of the amount of both fibrils and soluble aSyn species in each PFF preparation. This can be achieved by quantifying 1) the amount of monomers before initiating the fibrillization reaction; 2) the amount of aSyn converted to fibrils (Fig. 2A scheme, pellet) and the remaining aSyn soluble species at the end of fibrillization (Fig. 2A scheme, supernatant) using SDS-PAGE analysis. After 5 days (Fig. 2C), we typically see > 90-95% conversion of aSyn monomers to fibrils, as evidenced by the almost total absence of an aSyn band in the supernatant lane for the fibril batches 1-4. Fig. 2D shows the quantitative analysis of the extent of fibril formation from 12 independent fibril preparation experiments, estimated through SDS-PAGE band intensities. Two of the fibril preparations, batch 5 and 6, show higher levels of soluble aSyn species (batch 5: 12.3% and batch 6: 19.5%), which could indicate either incomplete fibrillization or the formation of less stable fibrils. Importantly, such variations are not captured by the standard qualitative analysis of fibril preparation using techniques such as EM and CD. Fig. 2E shows the EM images from three of the fibril batches which reveal predominantly straight and long fibrils. Thus, aSyn monomers are not detectable by EM and when present as minor species, their existence and levels can not be easily discerned by CD. Finally, it is important to emphasize that the fibrils exists in equilibrium with monomers and further manipulation of the fibrils could influence this equilibrium. For example, prior to their use in seeding-based assay, PFFs are subjected to sonication to induce the fibril fragmentation and to generate homogeneous (in terms of their average length, 50-150 nm) PFF resuspensions that are easily taken up by cells (Fig. 2B). Fig. 2F and G show an example of the average fibril lengths before and after sonication for a typical PFF preparation based on EM analysis. For the great majority of seeding-studies, only the average length and morphology of the fibrils are usually assessed. We strongly recommend that the amount of soluble aSyn species should also be assessed prior to and after sonication and preferably on the day the PFF preparations are used.

**Fig. 2:**
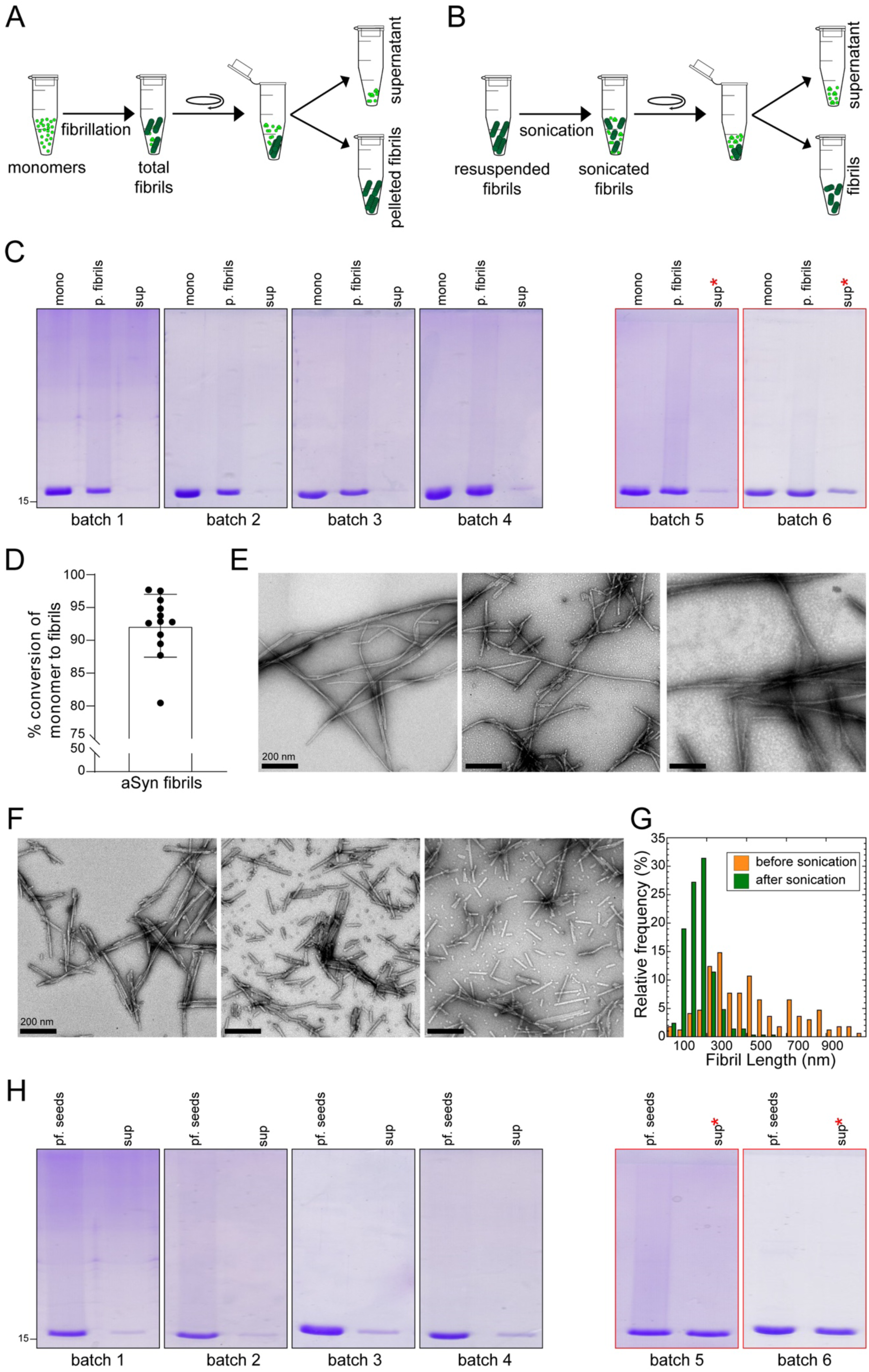
Schematic illustrations of (A) the separation of aSyn fibrils from a crude mixture of total aSyn fibrils following fibrillization and (B) following sonication (20% amplitude, one-second ON/OFF pulse cycle for 20 seconds), to assess the amounts of monomers/oligomers present in the supernatant fraction. C) SDS-PAGE analysis of the different batches of aSyn fibrils during fibrillization; mono: aSyn monomers prior to initiating fibrillization; p. fibrils: pelleted fibrils after resuspension; and sup: supernatant aSyn species following centrifugation at the end of fibrillization. D) Bar chart displaying the percentage of aSyn monomers converted to fibrils in different batches of fibril preparation. Bullet points represent the different batches of fibril preparation (n=15). E) EM images of total aSyn fibrils at the end of fibrillization showing three representative images from three random batches of fibril preparations. F) EM images of sonicated aSyn fibrils showing three representative images from three random batches. G) Differences in the lengths of the fibrils before and after sonication. H) SDS-PAGE analysis of different batches of aSyn fibrils following sonication; pf. seeds: the fibril seeds sedimented as pellets after sonication; sup: the soluble supernatant aSyn species following centrifugation. * in C and H denotes the increase in the amount of soluble aSyn species in the supernatant fractions in some non-reproducible fibril batches.

To illustrate how this could be done, the amounts of soluble aSyn species released after sonication, the sonicated fibril sample shown in Fig. 2B were subjected to ultracentrifugation at 100000*g* for 30 minutes, as described above. SDS-PAGE analysis showed the differences between the levels of sedimented fibril seeds and soluble aSyn species released after sonication for different PFF batches (Fig. 2H). For stable fibril preparations, we typically see ∼ 5-10% soluble aSyn after sonication (Fig. 2 H, batches 1-4). The stability of fibrils here is defined by their propensity to disassociate and the amount of soluble aSyn species detected after sonication. Depending on the stability of the fibrils, the amount of soluble aSyn released after sonication could vary significantly. For instance, batches 5 and 6 show approximately more than 30% soluble aSyn species in the supernatant generated by sonication.

Modifying the sonication parameters such as the length of sonication time (0-5-15-30-60 or 180 seconds) has been shown to strongly influence the length of the fibrils in the PFF preparations (Tarutani *et al*. 2016). We have also observed that variations in these parameters strongly influence the extent of soluble aSyn species released from the fibrils. When using handheld sonicators, the position of the sonicator with respect to both the sample and the wall of the tube were found to influence the release of soluble aSyn species from the fibrils. Therefore, we highly recommend exercising extreme care when performing this procedure. In addition, importance should be given to the determination of the ratio of soluble-to-fibrillar aSyn species, and to the assessment of the fibril length following sonication. Attention to these details is crucial for improving the reproducibility of seeding and aggregation assays and experiments.

### Detection and quantification of aSyn oligomers in PFF preparations

While the protocol described above could be used to quantify the levels of aSyn monomers and fibrils before and after sonication of each preparation, it is not suitable for quantifying oligomers in these preparations. Given the potent toxicity of aSyn oligomers (Fusco *et al*. 2017; Danzer *et al*. 2009; Mahul-Mellier *et al*. 2015), the presence of small amounts of oligomers could be a major contributor to experimental variability. In the following sections, we describe the extension of the protocol from detailed above to allow for the separation and quantification of oligomers from different aSyn samples, including PFF preparations.

**Table 2.**
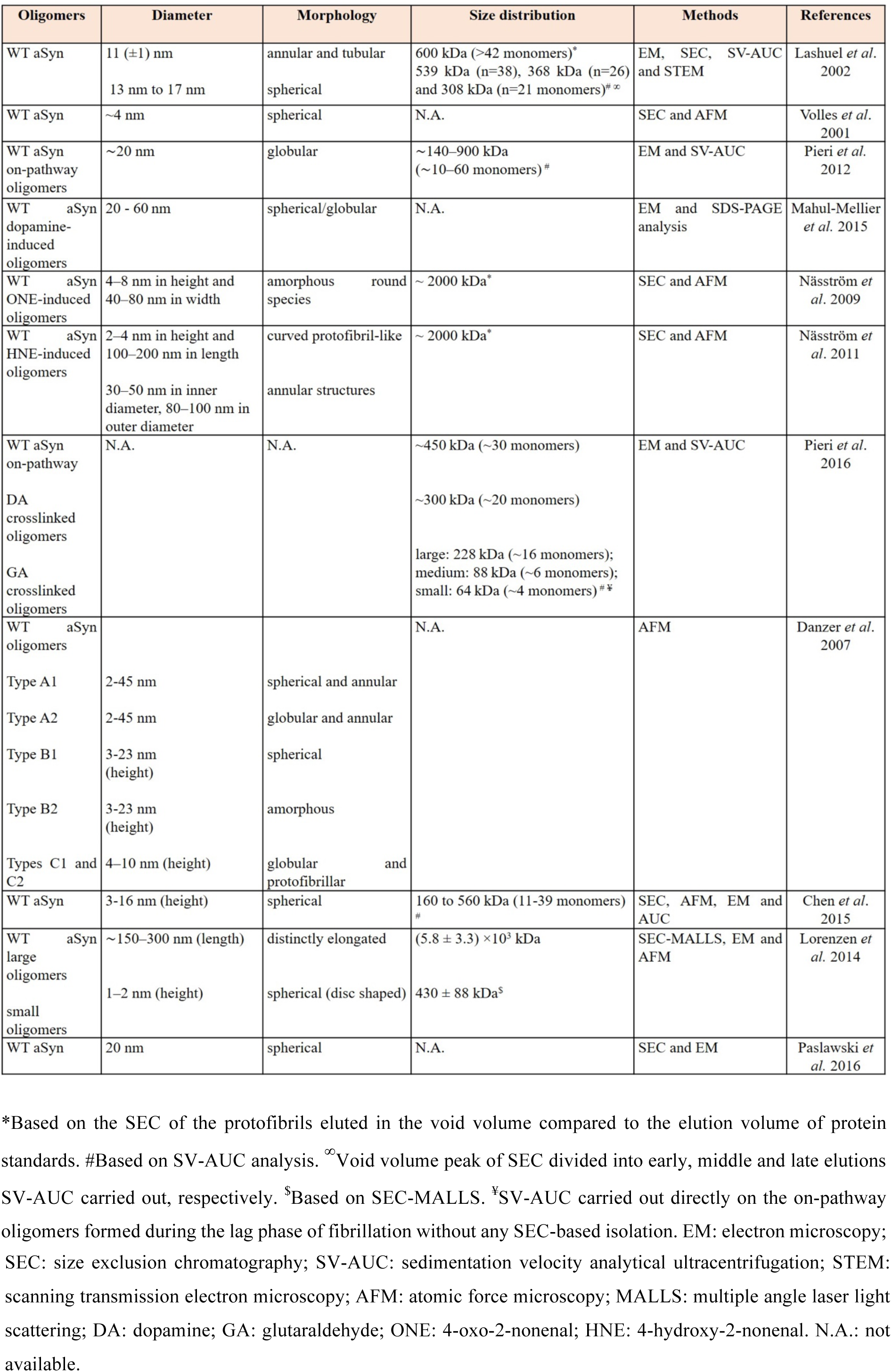
Summary of different protocols for the preparation of aSyn oligomers and their structural properties.

### Separation of monomers from oligomers

Despite the fact that different protocols for the preparation of aSyn oligomers have been developed (Table 2), analysis of the size distribution of the oligomers obtained using these preparations suggests that it is still possible to develop a single simple protocol that allows for the separation of monomers from oligomers (greater than a dimer). Based on the method of preparation, oligomers generally show different shapes (such as spherical, chain-like and annular structures), a wide range of sizes (usually ranging from 2-60 nm in diameter), and a molecular weight distribution ranging from 88-5800 kDa (Table 2). The large difference in the size and molecular weight between monomers and oligomers suggests that spin filters with a MWCO of 30 kDa or greater retain the oligomers on the membrane while allowing the monomers to pass through. However, when dealing with natively unfolded proteins such as aSyn, one must consider that a higher hydrodynamic radius exists, which makes aSyn behave as a globular protein of a larger size. For example, the aSyn monomer, which has a theoretical molecular weight of 14 kDa, behaves like a 57-60 kDa protein in size exclusion chromatography (Fauvet *et al*. 2012; Weinreb *et al*. 1996).

To determine which MWCO membrane provides the most efficient separation of aSyn monomers from oligomers, we first assessed and compared monomer recovery using 30 kDa, 50 kDa and 100 kDa MWCO membranes (Fig. 3A). As shown in Fig.3B, a complete recovery of aSyn monomers into the filtrate fraction was observed only when using a 100 kDa membrane and not a 50 kDa or a 30 kDa membrane. This is consistent with the higher predicted molecular weight of aSyn based on SEC (∼ 57 kDa). As the 100 kDa MWCO membrane was made from regenerated cellulose (Microcon), we sought to assess whether the nature of the membrane influences the recovery of the monomers. To that end, we compared aSyn monomer recovery through 100 kDa MWCO membranes made from regenerated cellulose (Microcon, used above) to that made from polyethersulfone (Vivaspin) (Fig. 3C). As shown in Fig. 3C, the efficiency of aSyn monomer recovery varied depending on the membrane composition. While regenerated cellulose membranes (Microcon) enabled the complete recovery of the monomers into the filtrate fraction, the polyethersulfone membrane (Vivaspin) retained approximately 20% of the protein in the retentate fractions. This analysis underscores the critical importance of evaluating the recovery efficiency of aSyn proteins prior to the use of spin filters to separate aSyn species or produce aggregate-free monomeric preparations.

**Fig. 3:**
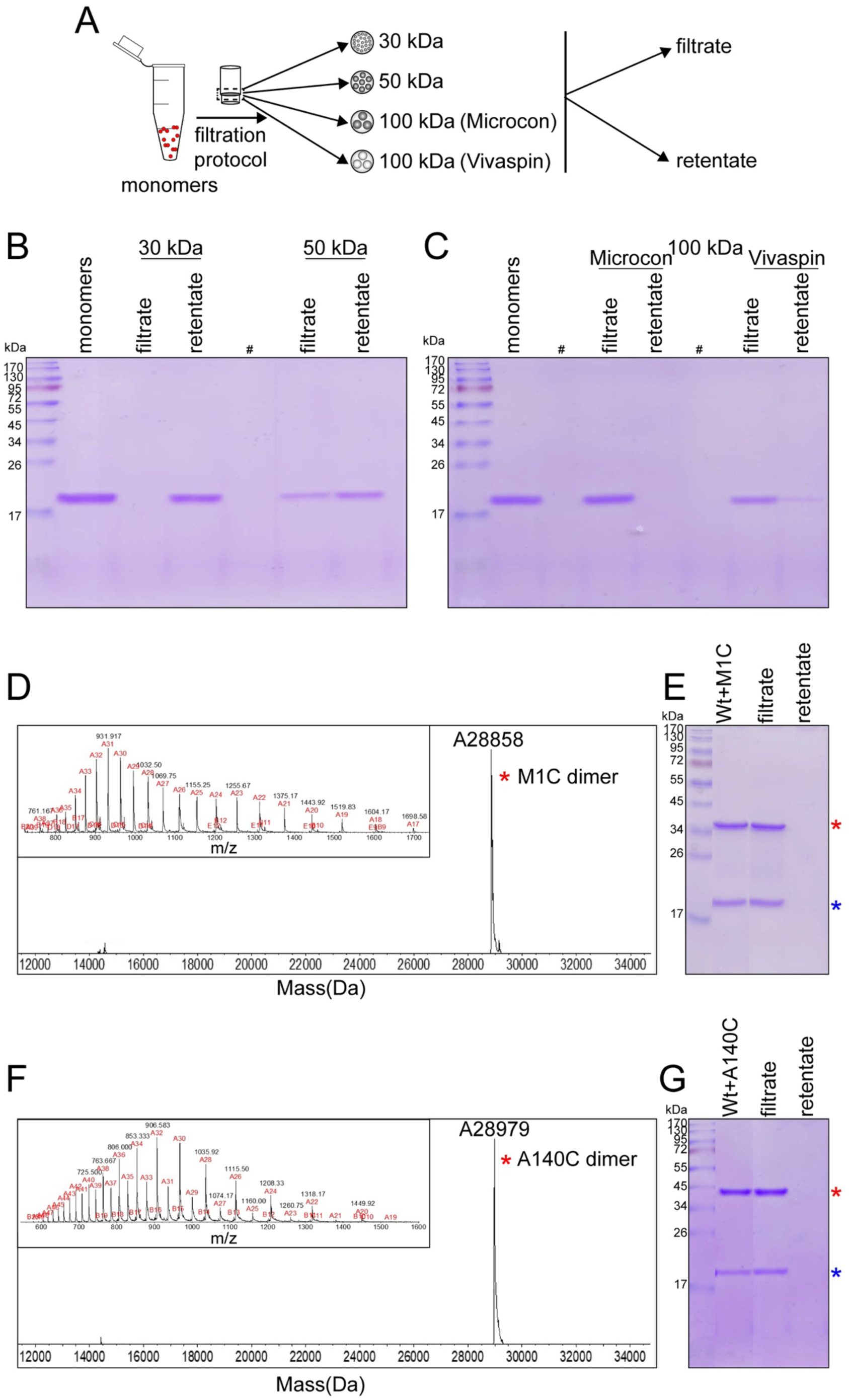
A) Schematic depiction showing the experimental setup used to assess aSyn monomer recovery using three different MWCO membranes - 30 kDa (Microcon), 50 kDa (Microcon) and 100 kDa (from commercial sources, Microcon and Vivaspin). B) SDS-PAGE analysis of the filtrate and retentate samples to assess the efficiency of the recovery of aSyn monomers after passing through 30 kDa, 50 kDa and 100 kDa membranes. C) Assessment of the recovery of aSyn monomers after filteration through a 100 kDa membrane from two different commercial resources, Microcon and Vivaspin. D) and F) Mass spectrometry analysis of M1C-linked (D) and A140C-linked (F) aSyn homodimers. E) and G) Assessment of the efficiency of recovery of equimolar mixtures of WT aSyn monomers and disulfide-linked aSyn dimers, M1C (E) or A140C (G), after filteration through aa to 100 kDa membrane (Microcon). Blue * denotes the WT aSyn monomers and red * denotes the dimeric forms of aSyn cysteine variant M1C and A140C.

Few studies have shown the existence of aSyn dimers in dynamic equilibrium with aSyn monomers (Marmolino *et al*. 2016; Coelho-Cerqueira *et al*. 2013). Although aSyn dimers have not been isolated as stable species, stable SDS-resistant dimers have been observed under conditions of oxidative stress that induces covalent chemical modification, for instance dityrosine crosslinking, and in the presence of polyphenol-based small molecules (Souza *et al*. 2000; Masuda *et al*. 2006; Hashimoto *et al*. 1999). Therefore, we assessed whether the dimeric forms of aSyn would pass through the 100 kDa MWCO filters. Towards this goal, we generated stable disulfide-linked aSyn homodimers using recombinant aSyn proteins bearing a cysteine residue at the N- or C-terminus of the protein (M1C and A140C, Fig. 3D and F). To determine whether the 100 kDa MWCO spin filters could separate monomers from dimers, an equimolar concentration mixture of WT aSyn monomers and homodimers was prepared and subjected to the same filtration conditions. Irrespective of the position of the cysteine mutation on the dimer (M1C or A140C homodimers), both monomers and dimers were recovered 100% in the filtrate fractions (Fig. 3E and G), and no traces of the protein were observed in the retentate fractions.

Next, we assessed whether the same protocol and membrane could be used to efficiently separate aSyn oligomers from monomers or isolate oligomers from a complex mixture of aSyn species. For this purpose, we generated aSyn oligomeric preparations using the protocol based on size exclusion chromatography purification, originally developed by Lashuel *et al*. 2002. EM analysis of the oligomer preparations before applying the filtration protocol shows the appearance of annular, tubular and spherical morphologies (Fig. 4 A) with diameters ranging between 6-14 nm (Fig. 4D). Previous sedimentation velocity and SEC studies have shown that these oligomeric preparations exhibit a size distribution with an average relative molecular mass that is slightly greater than 600 kDa (Lashuel *et al*. 2002).

**Fig. 4:**
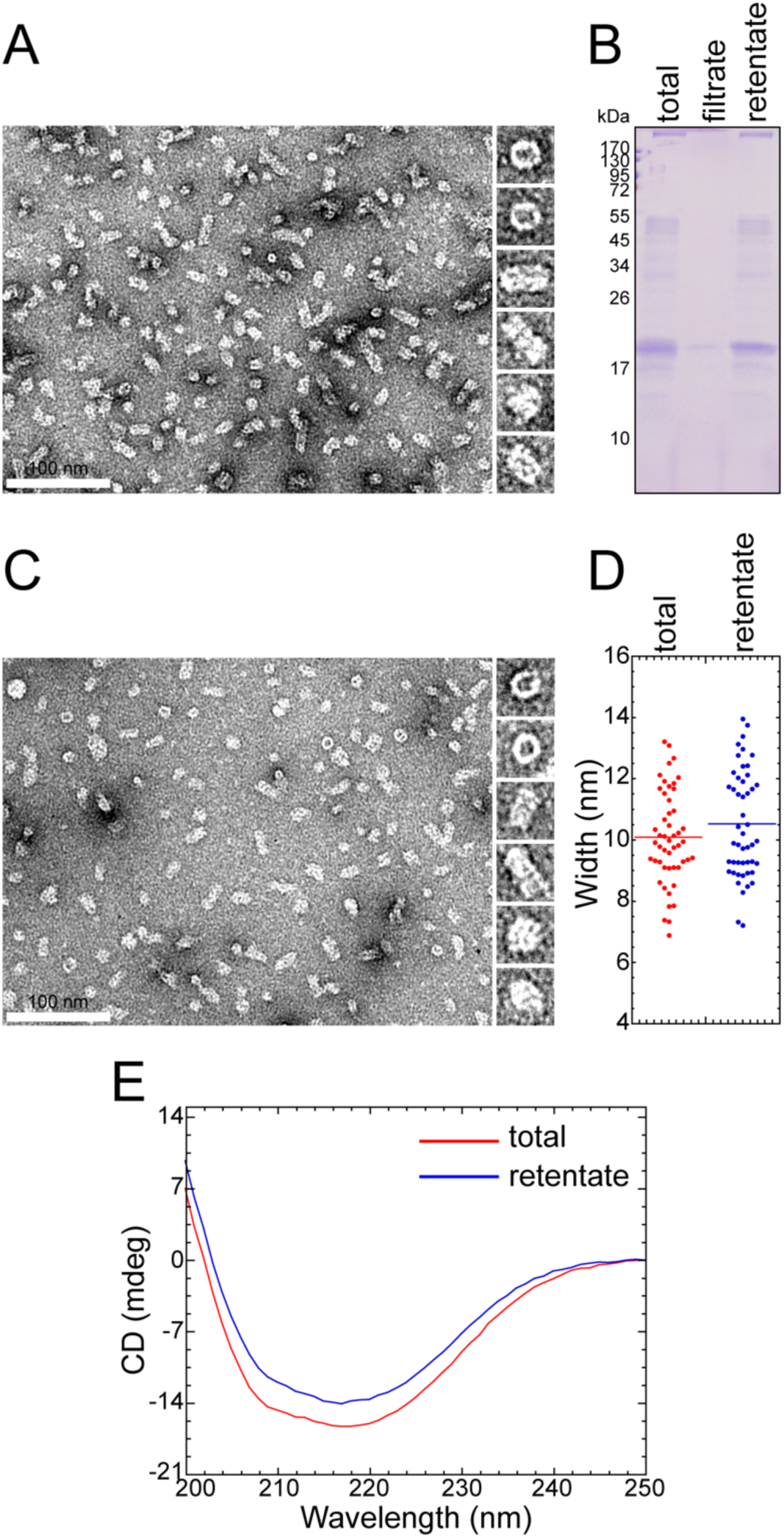
aSyn oligomers recovered through 100 kDa MWCO filtration did not alter their secondary structure, size or morphology. A) EM image of total aSyn oligomers used for the filtration protocol and the montage showing the most represented oligomeric structures. B) SDS-PAGE analysis of the total, retentate and filtrate samples of aSyn oligomers. C) EM image of retentate aSyn oligomers recovered after the filtration protocol and the montage showing the most represented oligomeric structures. D) Width analysis of the total and retentate recovered oligomers. E) CD spectra of total and retentate recovered oligomers.

As expected, based on their size and dimensions, the aSyn oligomers did not pass through the 100 kDa MWCO filters and were retained and recovered as the retentate. The filtrate fraction showed only a negligible trace of aSyn monomers. More importantly, EM analysis of the recovered oligomers revealed no significant changes in the size, and morphology distribution of the aSyn oligomers (Fig. 4C). Comparison of the CD analysis of the original oligomer samples and the oligomers recovered in the retentate also showed no changes in their secondary structures. These results show that filtration through 100 kDa MWCO filters allows for the efficient separation and recovery of aSyn oligomers and monomers and enables the rapid generation of highly pure samples of each species from samples containing a mixture of both species. Several studies have shown that using monomeric aSyn preparations that are free of any preformed fibrils or oligomers are essential to obtain reproducible and reliable aggregation kinetic profiles and results (Fredenburg *et al*. 2007; Singh *et al*. 2013; Buell *et al*. 2014). This is commonly achieved using SEC. However, this procedure requires at least 2-5 hours and is not suitable for experiments that involve comparative analysis of multiple samples or the use of highly precious aSyn species, e.g., site-specifically modified synthetic or semisynthetic proteins. Here, we show that a simple protocol based on filtration through a 100 kDa MWCO membrane provides a means for generating an aggregate-free aSyn preparation and the rapid removal or isolation of aSyn oligomers (> aSyn dimers).

### Analysis and separation of aSyn mixtures containing monomers, oligomers and fibrils

To demonstrate the robustness of the protocol, we tested it on an aSyn sample mixture composed of known concentrations (determined by UV absorbance) of monomers, oligomers and fibrils (referred to as the aSyn mixture), as illustrated in Fig. 5A. This mixture was then subjected to a centrifugation-based filtration protocol, and the aSyn species isolated in each step of the protocol were characterized using SDS-PAGE analysis, amino acid analysis, CD and EM. Fig. 5B shows the SDS-PAGE analysis of the mixture as well as aSyn samples recovered from each step of the protocol, i.e., the pellet (fibrils), supernatant (monomers and oligomers), filtrate (monomers) and retentate (oligomers). This analysis shows that we are able to recover each species. Fig. 5C shows the initial concentrations (based on the UV-absorbance readings) of individual aSyn samples (fibrils, oligomers and monomers) used in the aSyn mixture, and the concentrations of the recovered aSyn species isolated during the different steps of the protocol (determined by amino acid analysis). Comparison of the protein concentration estimation reveals that there is almost total recovery of the oligomers (10 μM was the initial concentration and 9.8 μM was the recovered concentration) and monomers (20 μM was the initial concentration and 17.6 μM was the recovered concentration), confirming that there was no significant reduction in the concentrations of soluble aSyn species during the recovery. The concentration of fibrils recovered (23 µM) was also similar (within error) to that of initial concentration (20 μM) added to the mixture.

**Fig. 5:**
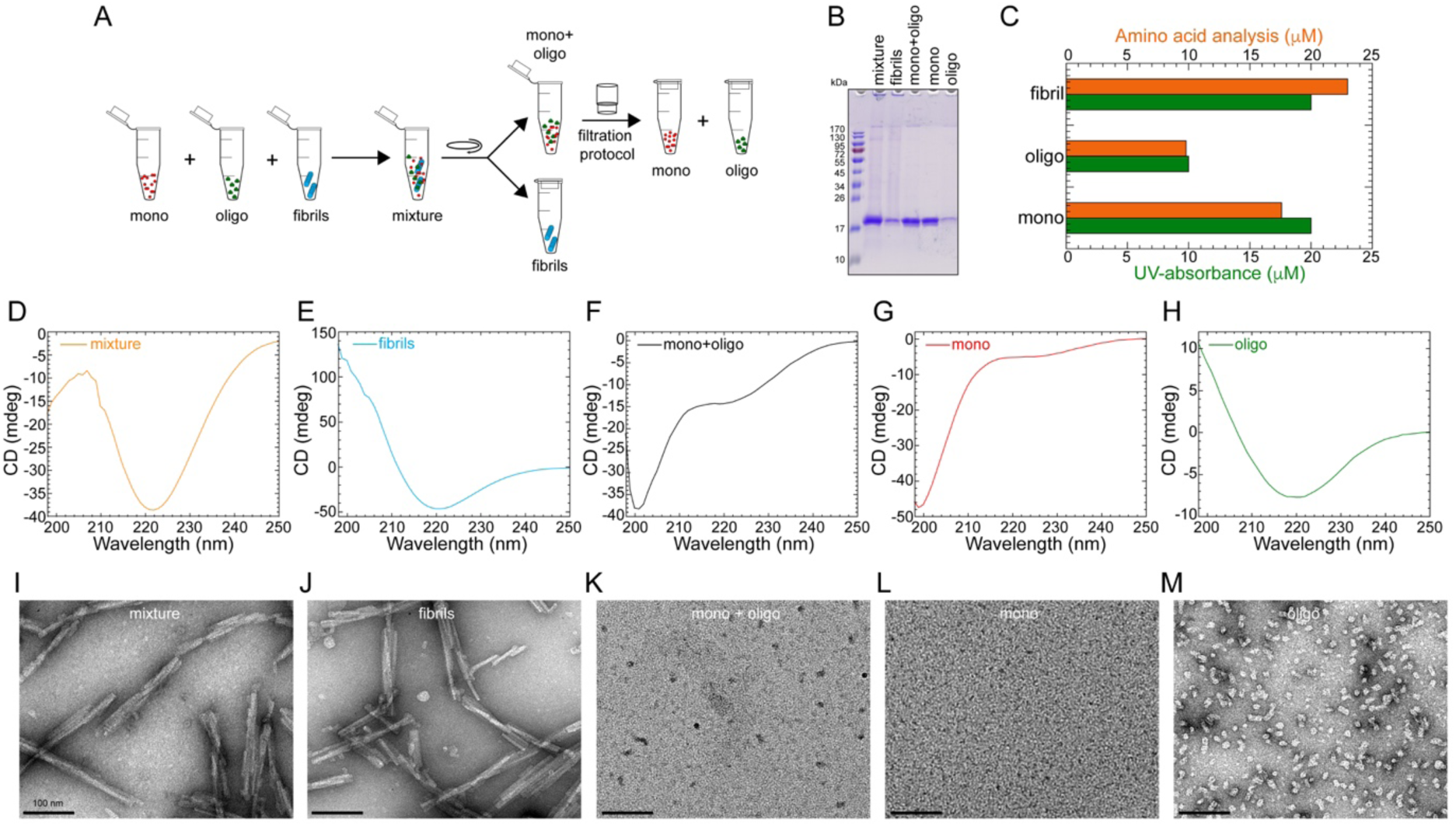
A) A scheme illustrating the experimental setup for assessing efficiency to separate aSyn species from a prepared mixture containing known concentration of aSyn monomers, oligomers and fibril. B) SDS-PAGE analysis of the samples collected at each step of the protocol. C) Comparison of protein concentration estimation (μM) of aSyn samples (fibrils, oligomers and monomers) from the initial concentrations, i.e., before the preparation of aSyn mixture (based on UV-absorbance) vs the obtained concentrations (based on amino acid analysis) of aSyn fractions isolated from the aSyn mixture during the steps of the protocol. D-H) CD spectra of the total aSyn mixture (D), fibrils (E), monomers and oligomers (mono+oligo) in the soluble aSyn supernatant (F), monomers (mono) in the filtrate (G) and oligomers (oligo) in the retentate (H) samples. I-M) Representative electron micrograph images of the aSyn mixture (I), fibrils (J), monomers and oligomers (mono+oligo) in the soluble aSyn supernatant (K), monomers (mono) in the filtrate (L) and oligomers (oligo) in the retentate (M) samples isolated from the different steps of the protocol.

The importance of this protocol is emphasized by the results obtained from CD and EM analyses. Fig. 5D and I show the CD and EM analyses of the total mixture of aSyn before subjection to the protocol (Fig. 5D and I) and the fibrils (Fig. 5E and J), monomers (Fig. 5G and L) and oligomers (Fig. 5H and M) after separation using the protocol. The EM image of the aSyn mixture is dominated by the fibrils (Fig. 5I). The oligomers are seen in the background and are often underestimated or overlooked in samples that are rich in fibrils, whereas the monomers cannot be detected by EM. The CD spectra are characterized by a minimum at 221 nm, which is consistent with a secondary structure that is dominated by *β*-sheets (Fig. 5D). This signal comes primarily from the oligomers and fibrils, both of which are rich in *β*-sheet structures. The relative contribution of the monomers to the CD spectra is not easy to delineate. This illustrates the difficulties in assessing the complexity and heterogeneity of aSyn samples containing multiple species by EM or CD, and underscores the critical importance of using protocols that provide a quantitative assessment of the distribution of the different aSyn species. The data in Fig. 5E-H and J-M show how this is achievable using the protocol described here. Fig. 5 E and J show the CD spectra and EM images of the aSyn fibrils separated by ultracentrifugation of the aSyn mixture at 100000*g* for 30 min at 4°C (Fig. 5A). As expected, and due to their enriched cross-*β* structure, the CD spectra of the sedimented and resuspended fibrils show the classical signal for the *β*-sheet structure (minimum at 221 nm, Fig. 5E), and the EM image (Fig. 5J) shows the appearance of straight and twisted fibrils. Analysis of the supernatant fraction containing soluble aSyn species (monomers and oligomers) by CD spectroscopy revealed a spectrum (Fig. 5F) that is dominated by a minimum at 200 nm and a shoulder minima band at 223 nm, consistent with the presence of both disordered (monomers) and structured (oligomers) aSyn species. Analysis of this sample by EM shows the presence of oligomers, but their morphological and size diversity is challenging to discern (Fig. 5K). Upon separation of the oligomers from the monomers by filtration through a 100 kDa MWCO filter, the CD spectra for the isolated monomer shows a single minimum at 199 nm without a shoulder, indicating the effective removal of all oligomers, which was also further confirmed by the absence of any oligomers in the sample (Fig. 5G and L). The recovered oligomers were resuspended in the same volume as the supernatant. They exhibited a CD spectra with a broad minimum centered at 219 nm (Fig. 5H), consistent with the presence of mixed secondary structure contents dominated by *β*-sheet structures. EM analysis of the recovered oligomers allowed more easy assessment of their morphological diversity and revealed the presence of spherical, annular and tubular-shaped structures (Fig. 5M). Taken together, these data show that applying our protocol to aSyn samples could reveal their complexity, and that relying solely on CD or EM analyses may lead to incorrect conclusions about the heterogeneity of the sample. This could partially explain some of the variations and/or irreproducibility of experimental observations in the field.

### Assessing the effect of PFF lyophilization on distribution of aSyn species

Very often, fibril and oligomeric preparations of aSyn are shared and exchanged between different research groups. The samples are usually frozen and shipped, or as more recently, done by Proteos (https://www.michaeljfox.org/grant/external-validation-proteos-alpha-synuclein-preformed-fibrils) stored and shipped in freeze-dried form (lyophilized powder). Previous studies have shown that subjecting aSyn samples to freeze/thaw cycles induces/increases oligomer/soluble aSyn species formation (Mollenhauer *et al*. 2017; Polinski *et al*. 2018; Stephens *et al*. 2018). The extent to which these storage procedures alter the biophysical properties of the fibrils will most likely depend on the type of fibrils present and the number of freeze/thaw cycles. Therefore, it is crucial to reassess the properties of the samples after thawing or resolubilization.

Here, we applied the centrifugation-based filtration protocol to an aSyn fibril sample, which was lyophilized and resolubilized in water (referred to as lyophilized fibrils). Fig. 6A shows the illustration of the steps and analysis we performed on this sample. SDS-PAGE analysis (Fig. 6B) revealed that only half of the aSyn protein is present as the mature and sedimentable fully grown fibrils as identified in the pellet fraction. The CD and EM analyses of the lyophilized fibrils shows the presence of *β*-sheet secondary structures (Fig. 6C) and long and straight aSyn fibrils (Fig. 6G). Similar to Fig. 5, aSyn monomers and oligomers are not easily detectable or quantifiable by these techniques. However, once we applied our protocol, one can clearly see that this sample of fibrils indeed contained a large fraction of monomers and a very small percentage of aSyn oligomers (Fig. 6B, E, F and I). SDS-PAGE analysis showed that approximately 5% of the soluble aSyn sample was retained in the spin filter recovered as the oligomer fraction. Filtered aSyn monomers show the typical CD spectra and characteristics of the unstructured aSyn species with a minimum signal at 200 nm (Fig. 6E). Again, CD and EM analyses of the soluble aSyn failed to detect the small percentage of existing oligomers, showing an bsence of any oligomers-like structures (Fig. 6H) and CD spectra displaying the absence of structured aSyn species. However, analysis of the recovered oligomers clearly showed the presence of small spherical shaped aSyn oligomers with an average diameter of approximately 12 nm (Fig. 6I, inserts) and the CD spectra (Fig. 6F) shows a peak minimum at 218 nm, revealing the structured oligomeric aSyn species. Taken together, this example illustrated how applying the centrifugation-based filtration protocol enabled the separation of complex mixtures of aSyn aggregate, and facilitated the detection and characterization of low abundance species which cannot be easily recognized in the EM images and contribute minimially to the CD spectra of such complex mixtures.

**Fig. 6:**
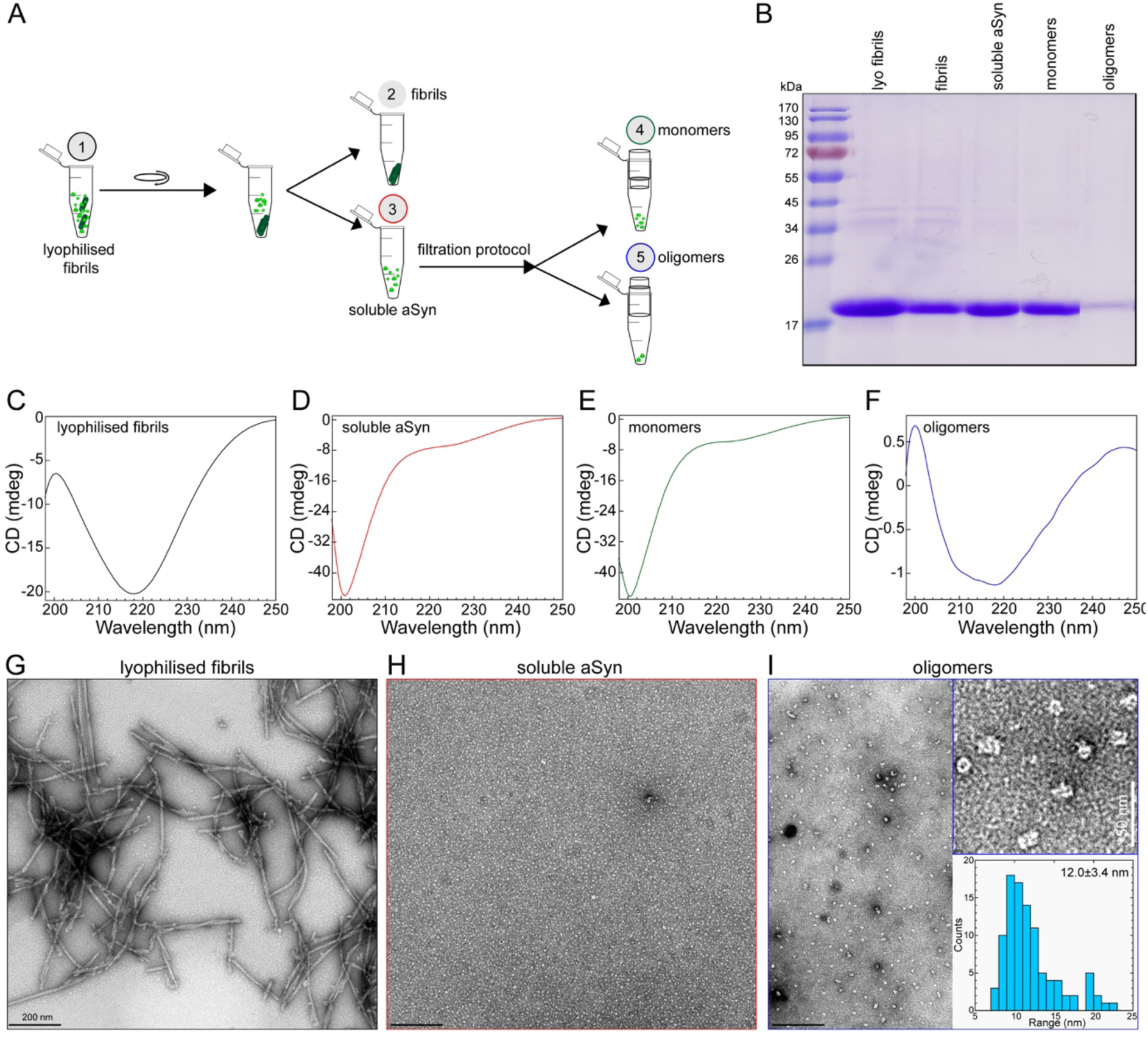
A) Schematic illustration showing the experimental setup of the centrifugation-based filtration analysis of lyophilized and resuspended aSyn fibrils (lyophilized fibrils). B) SDS-PAGE analysis of aSyn samples isolated from the different steps of the protocol. C-F) CD spectra of lyophilized fibrils (C), soluble aSyn (D), monomers (filtrate, E) and oligomers (retentate, F) in the samples. G-I) Representative EM images of lyophilized fibrils (G), soluble aSyn (H), and oligomers (retentate, I) in the samples from the different steps of the protocol. The top insert on I is the zoom of the EM images of the oligomer sample. The bottom insert of I shows the distribution of the diameter of oligomeric particles based on the quantification from EM images.

### Assessing the effects of sonication on fibril stability and disassociation

Another potential application we envisioned for this protocol was to assess the amount of monomers/oligomers released during the preparation of aSyn PFFs for *in vitro* and *in vivo* seeding aggregation studies. Between laboratories, aSyn fibrils are transported in the frozen/freeze-dried form and are sonicated before their application for seeding studies. We have examined the effect of sonication on different batches of freshly prepared fibrils and revealed its varying effects on the release of soluble species. (Fig. 2F-H). Here, we investigated the same on the stability of lyophilized fibrils.

The same sample of lyophilized and resuspended aSyn fibril used in Fig. 6 was used in this experiment, but the centrifugation-based filtration protocol was applied following the sonication of fibrils, as illustrated in Fig. 7A. The SDS-PAGE analysis shown in Fig. 7 reveals the differences in the amount of aSyn species recovered in each step of the protocol. Fig. 7G and H show the EM images of the lyophilized aSyn fibrils, which were long and straight (Fig. 7G) and similar to that observed in the Fig. 6G; however, sonication produced fragmented fibrils with length distribution ranging from 100-200 nm. The sonicated fibrils (Fig. 7C) retain their *β*-sheet rich structure. As expected, the soluble aSyn species in the supernatant after centrifugation exhibited a CD spectra that reflects a predominantly disordered conformations (Fig. 7D). aSyn oligomers were not easily visible by EM (Fig. 7I). However, upon filtration of the supernatant, we again observed nice separation of the unstructured monomers in the filtrate fraction (Fig. 7E) from the soluble oligomers with *β*-sheet structures in the retentate fraction (Fig. 7F). However, an important finding captured by the filtration protocol is that when compared to Fig. 6B, we see in Fig. 7B that there is an increase in the concentration of the the oligomeric sample recovered from the retentate fraction. This is also in agreement with the CD spectra which show a stronger CD signal (-3.8 CD (mdeg) on the y-axis, Fig. 7F) with a minimum at 218 nm compared to the weaker signal strength (at -1.2 CD (mdeg) on the y-axis, Fig. 6F), revealing an increase in the concentration of oligomers because of the effects of sonication on the lyophilized fibrils. Analogous to the oligomeric structures from Fig. 6I, the recovered aSyn sample (Fig. 7J) shows enrichment of spherical shaped oligomeric structures and the presence of very few short fragmented fibrils.

**Fig. 7:**
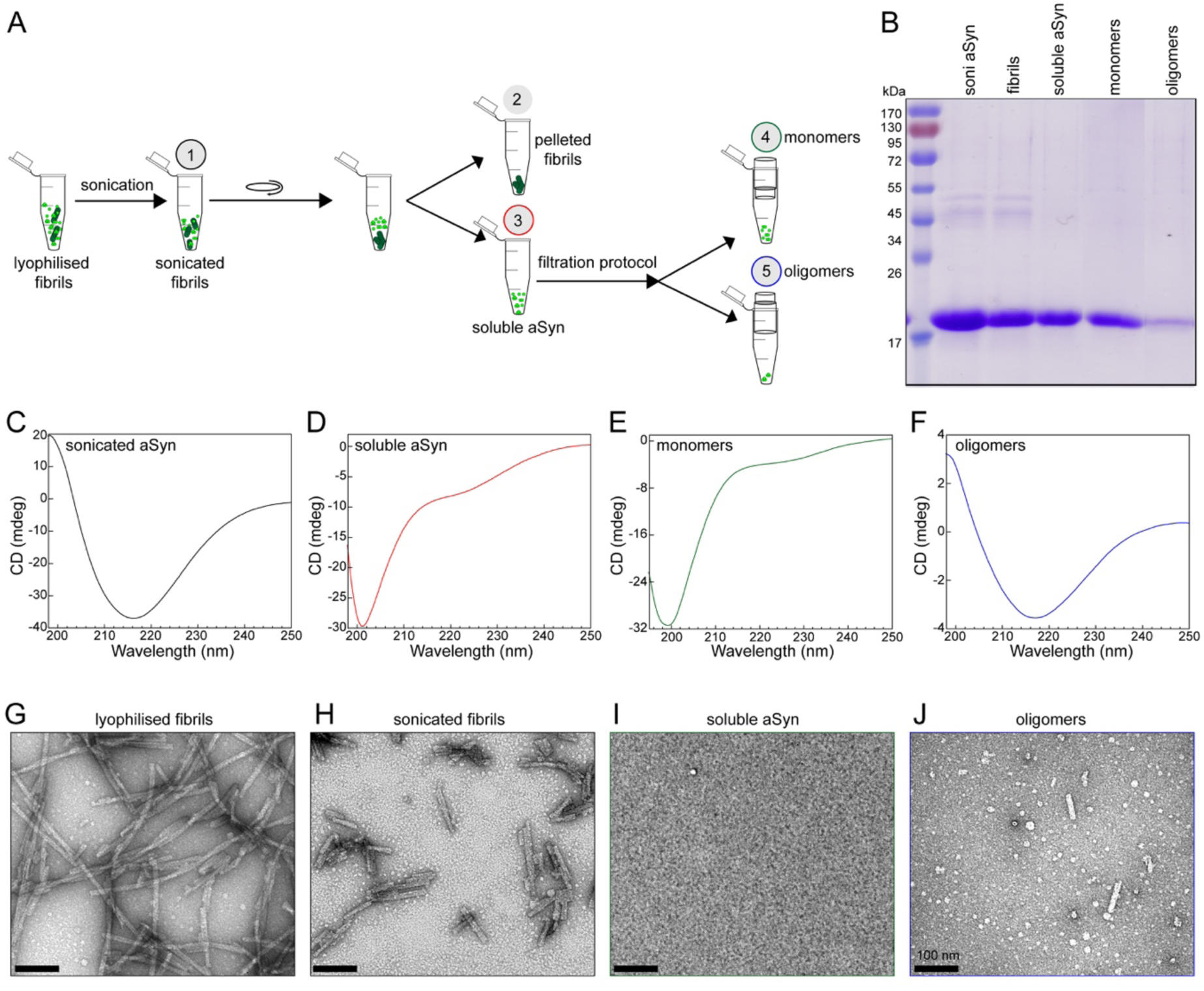
A) Schematic depiction of the centrifugation-based filtration protocol characterization steps following sonication (referred to as sonicated fibrils) of the lyophilized fibrils. B) SDS-PAGE analysis of aSyn samples isolated from the different steps of the protocol. C-F) CD spectra of sonicated fibrils (C), soluble aSyn (D), monomers (filtrate, E) and oligomers (retentate, F) in the samples. G-I) Representative electron micrograph images of lyophilized fibrils (G), sonicated fibrils (H), soluble aSyn (I), and oligomeric (retentate, J) samples.

In summary, the centrifugation-based protocol presented here is flexible, i.e., it can be easily applied to the same aSyn fibrils (Proteos) resuspended after lyophilization and following sonication. Very importantly, the protocol shows the robustness of separating the different aSyn species in the same aSyn sample, such as fibrils, monomers and oligomers, even when present at levels that are not detected by CD and EM analyses, and allows the study of each species carefully in isolation. Furthermore, the protocol captures changes in the concentration of aSyn oligomeric species following manipulation of the samples, e.g., PFF sonication. Such careful characterizations of the samples will improve the reproducibility of PFF preparations, which will lead to more reliable and reproducible results from experiments aimed at assessing the structural properties, interactome, toxicity, seeding capacity and pathology spreading of aSyn fibrils.

## Discussion

There is a consensus that the heterogeneity and variation in the distribution of different species (monomers, oligomers and fibrils) in aSyn aggregate preparations are the major factors contributing to the experimental differences and irreproducibility in aSyn research today. Previous efforts to address these experimental differences have focused on comparing the polymorphisms and structural properties of aSyn fibrils or oligomers as determined by TEM or AFM which do not allow for quantitative determination of the relative ratio of the different species in the sample. We believe that advancing our understanding on the role of aSyn aggregation in the pathogenesis of PD requires the development of reliable methods that not only enable the preparation of different species of aSyn or isolation of distinct aSyn species from complex sample mixtures but also quantitative assessment of the distribution of different aSyn species in such preparations. This is crucial since most aSyn oligomeric and fibrillar species are in dynamic equilibrium with monomers and could undergo conformational and quaternary structure changes in response to changes in their environment. Such methods should be accessible to most research laboratories and be suitable for the analysis of small amounts/volumes of samples. While several biophysical techniques have been widely used to characterize the size, morphology and structural properties of aSyn aggregates, those methods usually require high quantities of protein and special instrumentation and/or do not allow for the analysis of multiple samples within a short period of time (1-2 hours). These limitations make it very difficult to achieve detailed analysis of the distribution of aSyn samples prior to performing biophysical or biological studies and/or during these studies.

Herein, we describe a simple centrifugation-based filtration protocol that overcomes the aforementioned limitations. The protocol is based on multiple separations and analytical procedures that have been validated by multiple research groups. This protocol offers several advantages: 1) it requires only basic laboratory tools and equipment, which makes it easy to implement in most research laboratories, 2) it can be applied to very low sample volumes (50-500 μL); 3) it allows for isolation and characterization of each species at submicromolar concentrations and 4) it can be performed in a very short time (3-4 hours). To illustrate the utility of this protocol, we presented several examples that demonstrate how it can be used to: 1) assess the distribution of aSyn monomers, oligomers and fibrils in aSyn preparations; 2) isolate small amount of each of these species from complex mixtures; 3) investigate the extent of monomer and fibril release from aSyn fibrils; 4) capture the batch-to-batch variability in aSyn PFF preparations and changes in aSyn species upon manipulations of these preparations (e.g. lyophilization and sonication). We also showed that it enables the detection of small amount of oligomers in aSyn fibril preparations, the presence of which is usually undetectable or unappreciated by standard techniques used to characterize fibril preparations (e.g. CD and EM). These capabilities will contribute significantly to improve sample-to-sample variations and improve the level of characterization of monomeric, oligomeric and fibrillar aSyn samples used in structure-function/dysfunction relationship studies. This is crucial to allow accurate interpretation of experiments and comparison of results obtained using different preparations. This simple protocol has wide-ranging applications beyond what is described above. For example, it can be used to assess the disaggregase activity of small molecules and chaperones, allow determination of amounts of fibrils remaining or monomers release upon co-incubation with chaperones or drugs, and allow insight into the mechanism by which these disaggregases disassemble the fibrils, i.e., through fibril fragmentation, oligomer formation or monomer release. It is worth reminding that this protocol has been developed and validated for the characterization of aSyn samples in simple buffer solutions. We plan to explore the possibilities to modify it and extend its application to the characterization of aSyn species in complex samples derived from culture media, cell extracts, biological fluids or brain homogenates.

## Acknowledgement

We thank Somanath Jagannath for his assistance during the preparation of WT aSyn oligomers and Melek Firat Altay for her critical review of the manuscript. This project was funded by the École Polytechnique Fédérale de Lausanne and the Michael J Fox Foundation.

